# Using Machine Learning to Identify True Somatic Variants from Next-Generation Sequencing

**DOI:** 10.1101/670687

**Authors:** Chao Wu, Xiaonan Zhao, Mark Welsh, Kellianne Costello, Kajia Cao, Ahmad Abou Tayoun, Marilyn Li, Mahdi Sarmady

## Abstract

**Background:** Molecular profiling has become essential for tumor risk stratification and treatment selection. However, cancer genome complexity and technical artifacts make identification of real variants a challenge. Currently, clinical laboratories rely on manual screening, which is costly, subjective, and not scalable. Here we present a machine learning-based method to distinguish artifacts from *bona fide* Single Nucleotide Variants (SNVs) detected by NGS from tumor specimens.

**Methods:** A cohort of 11,278 SNVs identified through clinical sequencing of tumor specimens were collected and divided into training, validation, and test sets. Each SNV was manually inspected and labeled as either real or artifact as part of clinical laboratory workflow. A three-class (real, artifact and uncertain) model was developed on the training set, fine-tuned using the validation set, and then evaluated on the test set. Prediction intervals reflecting the certainty of the classifications were derived during the process to label “uncertain” variants.

**Results:** The optimized classifier demonstrated 100% specificity and 97% sensitivity over 5,587 SNVs of the test set. 1,252 out of 1,341 true positive variants were identified as real, 4,143 out of 4,246 false positive calls were deemed artifacts, while only 192(3.4%) SNVs were labeled as “uncertain” with zero misclassification between the true positives and artifacts in the test set.

**Conclusions:** We presented a computational classifier to identify variant artifacts detected from tumor sequencing. Overall, 96.6% of the SNVs received a definitive label and thus were exempt from manual review. This framework could improve quality and efficiency of variant review process in clinical labs.

## Introduction

A large number of unique and non-recurrent somatic and germline variants may exist in a cancer genome*(1)*. Clinical interpretation of these mutations is key for tumor stratification and subsequent treatment selections*(2)*. However, the diversity of somatic events that occur in heterogeneous tumor clones and technical artifacts make identification of *bona fide* genomic variants using next generation sequencing (NGS) technology a challenge*(3)*. Specifically, single nucleotide variants (SNVs) constitute the majority of the somatic variants of the cancer genome. These variants may only be present in a small portion of the sample DNA due to the subclonal events or contamination by normal cells*(4)*. The abundance of variant calls derived from inherently noisy NGS data, such as pseudogenes, sequencing artifacts or low coverage regions makes it even more arduous to identify the real somatic SNVs.

The choice of variant calling algorithms has a critical and direct impact on the outcome of the clinical laboratory findings, therefore the algorithms must demonstrate high robustness, sensitivity, and specificity. Many algorithms, such as VarScan*(5)*, SomaticSniper*(6)*, and MuTect*(4)*, incorporate unique models and varying information from the sequencing data, which leads to different performance characteristics. For instance, a highly sensitive algorithm is capable of detecting more real variants but may suffer from reporting higher rate of false positive calls *(7)*. Although lower specificity may be addressed through validation using an orthogonal method such as Sanger sequencing, it could be costly for clinical laboratories due to the high number of variants to be confirmed from a large sequencing panel *(8)*. Additionally, confirming somatic mutations with low allele fraction may be challenging *(9)*. A number of comparative studies have revealed the lack of concordance among different variant calling methods*(10, 11)*. To address this issue, some studies have suggested improved performance using ensemble or consensus approaches to detect somatic and germline variants *(12, 13)*.

While combining results from multiple variant callers increases sensitivity, it often yields large number of variants which pose a challenge for manual review and analysis in clinical labs. Due to the clinical demand for extremely high sensitivity and the complex nature of cancer genomes, noise, such as artifacts, may be introduced into the DNA sequencing datasets and can easily overwhelm the variant call sets*(14)*. A number of bioinformatics strategies have been proposed to perform variant refinement on the raw variant call set to remove likely false positives depending on caller-specific metrics such as mapping quality and strand bias*(11, 15, 16)*. These approaches apply a combination of filtration schemes on detected variants based on empirical observations, without systematically investigating the optimal cutoffs for each of the features to achieve the best performance. Further, clinical-grade sequencing and interpretation require additional quality-assurance methods to ensure the validity of the variants detected from the algorithms*(17)*. For instance, an in-house database of well-annotated variants is strongly recommended to characterize the mutations frequently encountered by the lab and hence facilitate this process*(2)*.

Quality control screenings are indispensable to filter sequencing artifacts and other non-reportable variants before assessing the clinical significance of the remaining variants. Visual inspections are commonly implemented in clinical laboratories for variant screening*(18, 19)*. A recent study has developed a deep learning-based approach to automate the variant screening process*(20)*. The computational models were trained on adult clinical tumor sequencing and public datasets, achieving high classification performance. Despite the wide collection of attributes and sophisticated methods, the optimized models did not achieve 100% sensitivity or specificity. Additionally, as the histopathological traits and molecular characteristics of pediatric tumors diverge from adult tumors *(21)*, the mutation landscape of pediatric tumors is drastically different from that of adult cancers *(22)*. Therefore, continued refinement of computational methods to improve variant review is necessary *(23)*.

Since SNVs constitute the majority of the detected variants in tumor samples*(21)*, and greater complexity of sequencing artifacts is observed in formalin-fixed paraffin-embedded (FFPE) tissues, we limited our study to SNVs of non-FFPE pediatric tumor samples*(24)*. In the following sections, we detail the design and assessment of the computational framework to automatically perform variants screening on pediatric tumor samples. We then demonstrate the optimized model can improve the accuracy and efficiency of tumor variant classification.

## Materials and Methods

### Sequencing and Clinical Bioinformatics Pipeline

Variant data sets used for this study were compiled from pediatric cancer patients who underwent molecular testing of hematological or solid tumor NGS targeted gene panels at the Children’s Hospital of Philadelphia (CHOP). The solid tumor panel comprised 238 genes while the hematological cancer panel comprised 118 genes *(25)*. For each of the clinical samples, regions of interest (ROI) were captured using Agilent SureSelect QXT target enrichment technology. FASTQ data generated by Illumina MiSeq/HiSeq sequencers was aligned to the hg19 reference genome using Novoalign*(26)*. The average coverage for the panels was 1500X with 99.7% of the ROI fully covered at equal to/greater than 100X. After alignment, four different variant callers were used to achieve a high detection sensitivity, including Mutect*(4)*, Scapel*(27)*, FreeBayes*(28)*, and VarScan2*(5)*. If a variant was detected by any of the tools, it was retained for downstream analysis.

### Manual Inspection

In the manual variant review process in the cancer diagnostics lab at CHOP, an SNV was deemed a sequencing artifact if at least two of the followings were true:

- High allele ratio (VAF) and could be visually seen in at least two normal controls of similar VAFs
- Low mapping quality
- High strand bias in both patient sample and control samples
- Supported by no more than two unique paired reads when the coverage at the locus was at least 50X
- Located in difficult genomic regions that were susceptible to potential PCR amplification errors, such as poly A/T regions, or of paralogous alignment quality *(29)*

Several healthy samples were selected to serve as negative controls to assist visual inspection. These samples were thoroughly investigated to be free of known pathogenic mutations of cancer genes in the panels and underwent same sequencing and bioinformatics processing as patient samples. The variant of interest was compared to the same genomic coordinate in these negative control samples in Integrative Genomics Viewer (IGV)*(30)*. The premise is if a variant under investigation could be observed in a similar manifestation in the control samples, it is likely called due to technical or algorithmic errors that universally affect other samples as well. Because of the complexity of cancer genomes and the nature of NGS, many thresholds in the criteria were empirically derived and refined over time. To mitigate the subjectivity introduced by personal bias, two independent reviews of the same variant were performed by different genome scientists, which made the procedure even more laborious and thus not scalable.

### Data Generation and Feature Selection

A total of 11,278 SNVs from 291 individual tumor samples of pediatric cancer patients from 9 cancer types and more than 30 subtypes of tumors (supplementary Figure 1) were compiled for the study. Each SNV was manually reviewed and labeled as either real (TP, including reportable variants, polymorphisms, intronic and synonymous variants) or non-reportable (FP, i.e. sequencing artifacts). Similar to previous machine learning applications in genomics *(31)*, data was randomly split into three subsets that were mutually exclusive: training, validation, and test sets. The training set comprised of 3,362 variants, from 61 solid and 23 hematology tumor specimens, respectively. The validation set comprised of 2,329 variants (32 solid/34 hematology tumor specimens), while the test set comprised of 5,587 variants (69 solid/72 hematology tumor specimens). The breakdown of the variants of these datasets was summarized in Figure 1.

**Figure 1.**
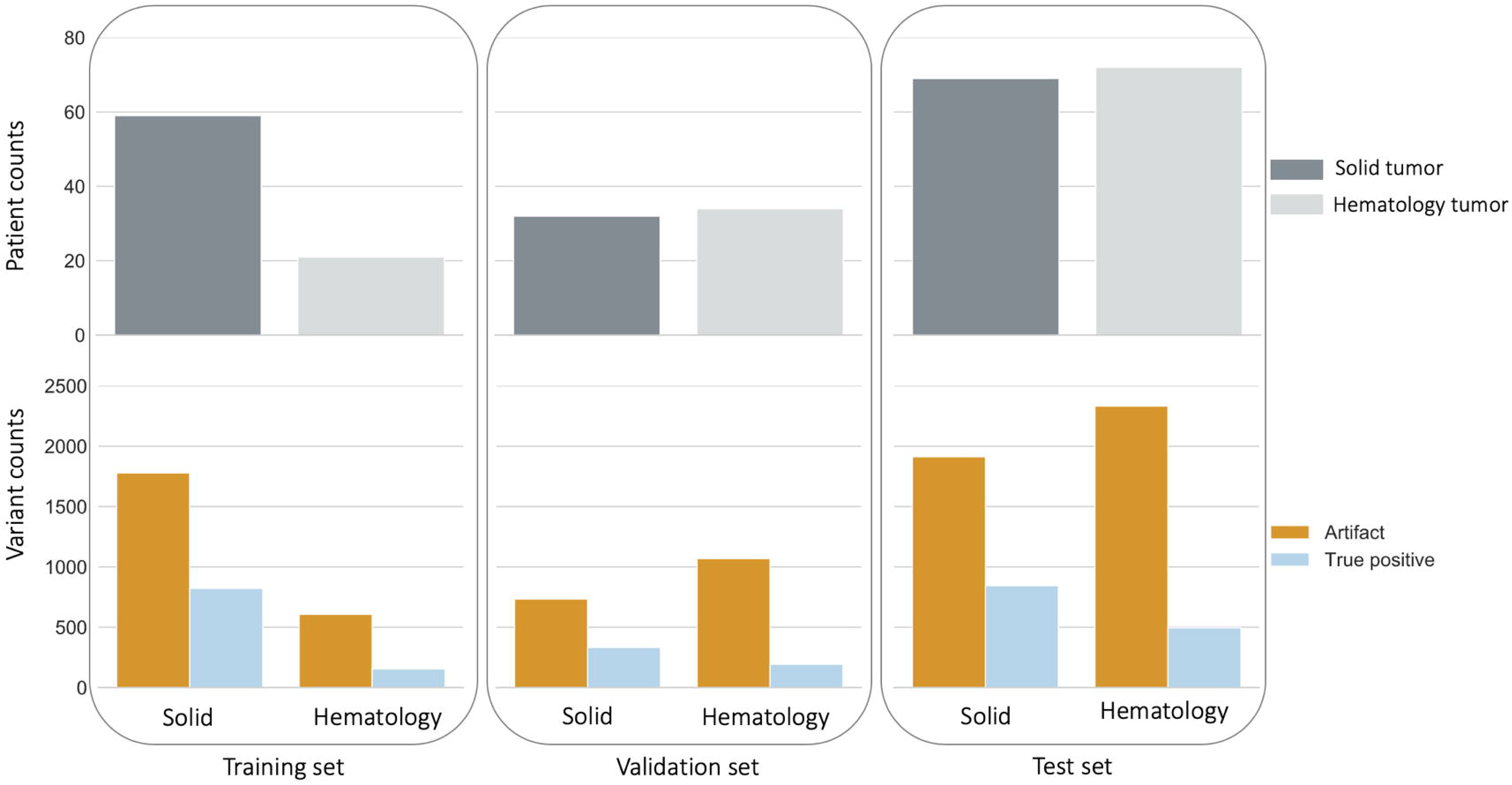
Variant data used in the study. The training set comprised 61 solid and 23 hematology tumor samples, including 976 true positive variants (821 solid/155 hematology tumors) and 2386 artifacts (1779 solid/607 hematology tumors) respectively. The validation set comprised 32 solid and 34 hematology tumor samples, including 526 true positive variants (333/193) and 1803 artifacts (734/1069) respectively. The test set comprised 69 solid and 72 hematology tumor samples, including 1341 true positive variants (845/496) and 4246 artifacts (1913/2333) respectively.

A pseudo-score based on the ENCODE mappability track *(32)* was derived to assess sequence uniqueness of each exon *(33)*. Variants from computationally inferred pseudo regions were marked in the clinical bioinformatics pipeline. These variants were challenging to review and were always confirmed by Sanger sequencing in case of clinical relevance, and hence were not included in the variant dataset of this study.

Guided by the manual inspection process, we started with a collection of attributes for each of the variant such as alternate allele coverage, minor allele fraction, etc. Univariate feature selection based on chi-squared test was performed to remove features that were less informative, such as mapping quality of the aligned reads. The following features were selected to represent each SNV in the computational model:

- Alternate coverage: number of unique reads supporting the alternate allele
- Strand bias: imbalance between aligned reads supporting the alternate allele on opposing strands, higher values indicated greater bias:

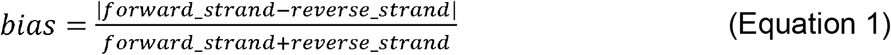
- Variant allele fraction (VAF): ratio between unique reads supporting the alternate allele and the total number of reads at the locus
- Dissimilarity to normal control samples: this feature captures the separation between the variant of interest and the characteristics of the alleles at the same genomic coordinate in normal control samples. A three-component vector was composed integrating alternate coverage, strand bias and VAF for the variant of interest and the same chromosomal locus on the normal control sample. The metric was measured by the Euclidean distance between the two vectors:

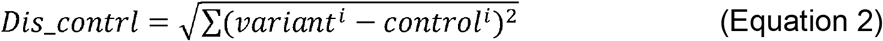

Two control samples with the highest VAF were selected to compare with the variant of interest using Equation 2, because the probability of both unrelated samples carrying the same somatic variants was extremely low.
- Batch effect: the metric indicated the separation between the variant of interest and the characteristics of the same genomic coordinate on the other samples processed in the same batch. One sample besides the patient sample from the batch exhibiting the highest VAF was selected to compare with the variant of interest using Equation 2.

To assess the separation of data in an unsupervised manner, a principal component analysis was performed on the data and the result suggested the two classes were largely separable using the selected features (Figure 2).

**Figure 2.**
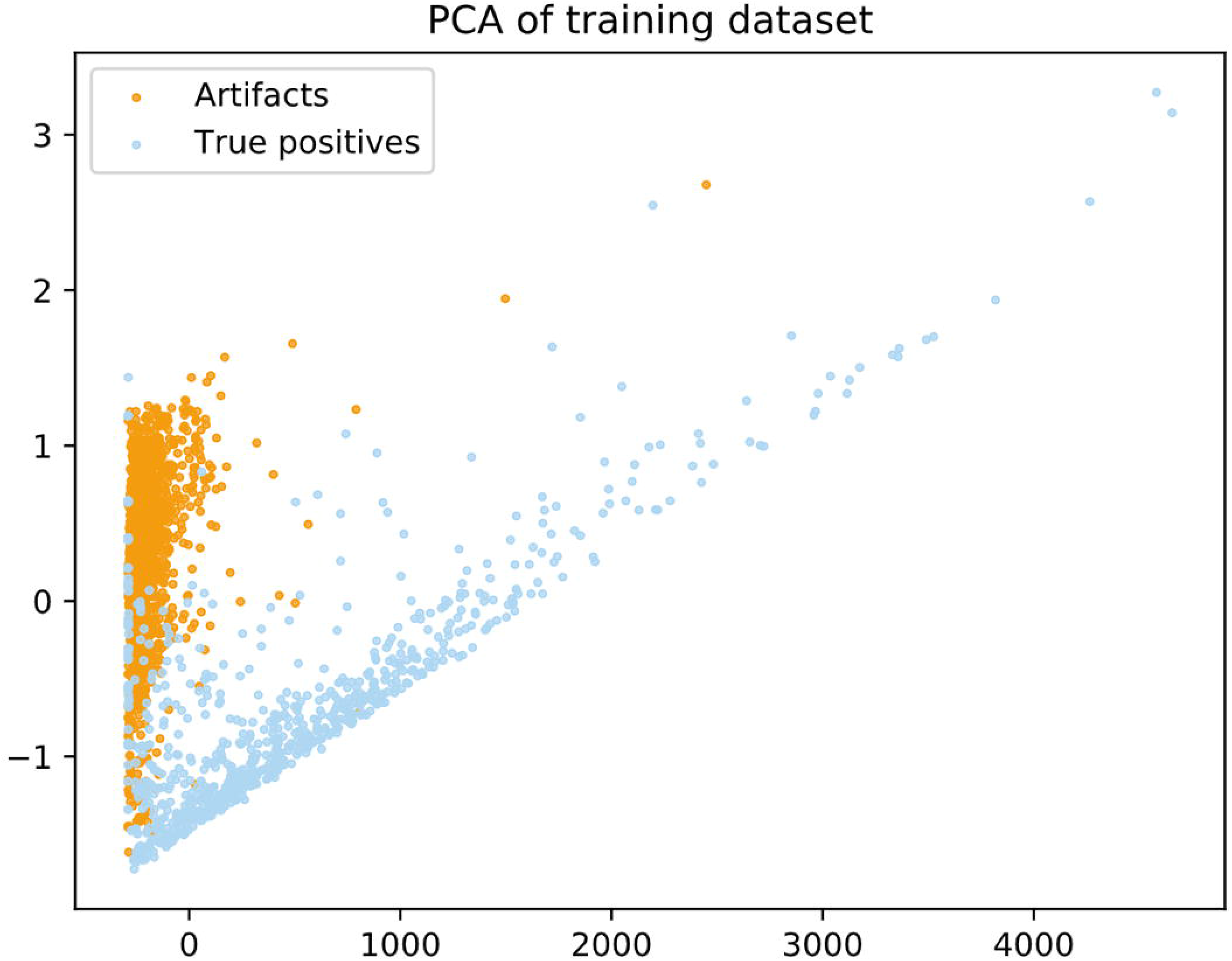
Variants of training data represented using the first two components from the Principal Component Analysis. The plot indicates the two classes are largely separable despite a small degree of overlapping.

### Computational Framework Training, Tuning and Testing

Random forest (RF) algorithm as an ensemble approach was implemented since it has been demonstrated to be adaptive to correlated features and prevent overfitting for genomic data *(34)*. The models were trained, validated, and tested using the Python *Scikit-learn* package *(35)*.

A proof-of-concept model was learned using the training set, which achieved 100% sensitivity and specificity with a 0.98 F1 score in 10-fold cross-validation on the two-class training set. Following this, fine-tuning parameters of the model was carried out by evaluating the performance using the validation set. To achieve clinical assurance, an imperative objective in this step was to derive a three-class classifier from the baseline model, for the third class being “uncertain”. Systematic errors may contribute to the ambiguity such as insensitive variant calling for variants with low VAF, low coverage or imperfect alignment. These errors can be difficult to analyze, as the source of errors could be lost in most existing evaluation methods. *(36)* Therefore, the third class may include variants of complex feature manifestations and require further manual inspection with additional information. During this process, different ranges of values over a set of parameters were evaluated, resulting in over 1,000 classifier instances, before an optimal model was identified. Following this, the optimal model was benchmarked on the independent test set. The overall workflow of this study is demonstrated in Figure 3. To reflect the higher cost of type 2 errors than type 1 errors in clinical laboratories, different weights were assigned to true and false outcomes as explained in Results.

**Figure 3.**
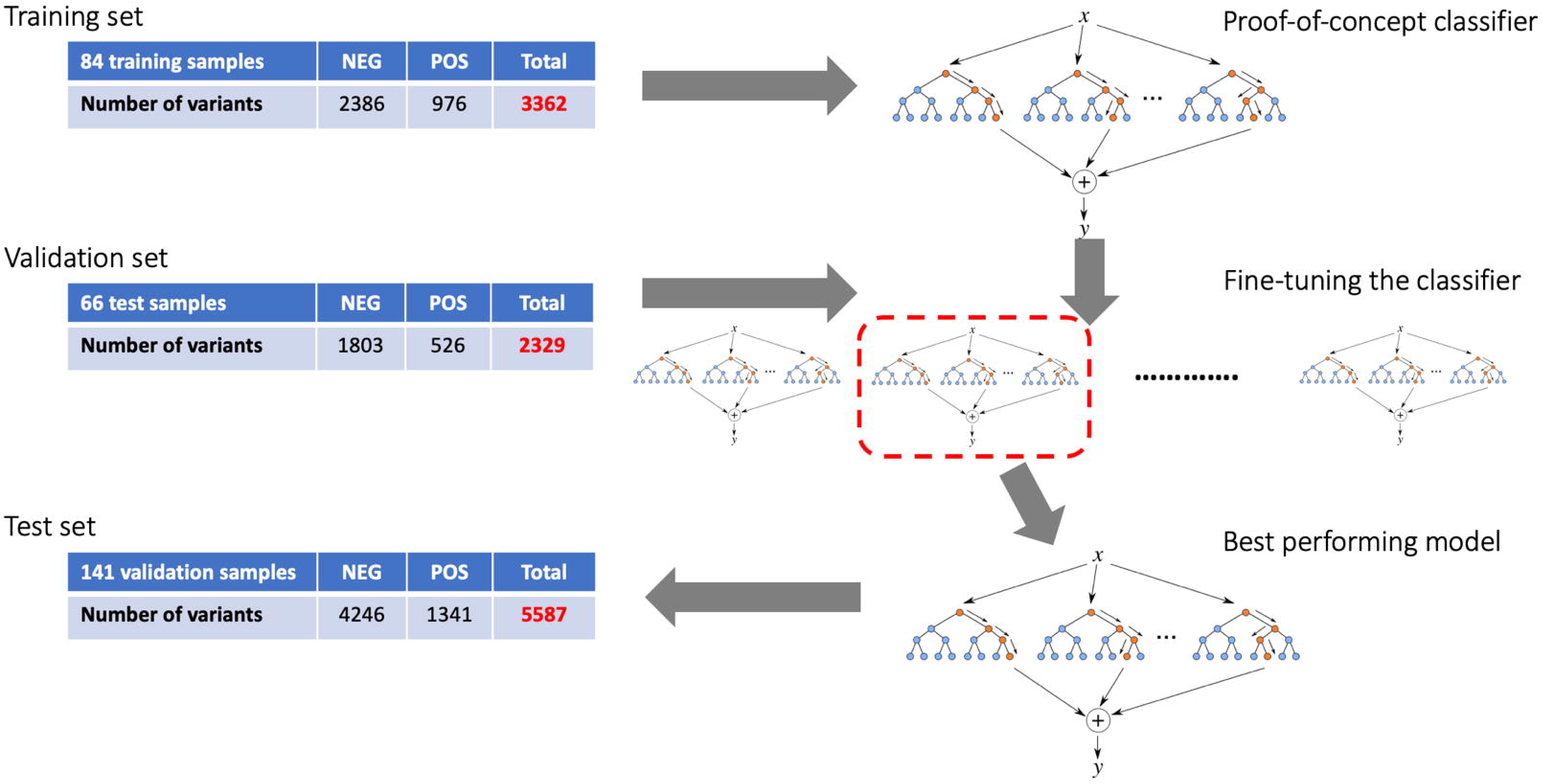
Workflow of computational classifier development. A baseline model was trained using the training set. Different sets of parameters were evaluated to identify the best performing model using an independent validation set, before benchmarked using the test set.

## Results

### Baseline Model from Training Set

The training set consisted of 3,362 labeled SNVs from 84 somatic tumor samples, and each SNV was represented by the features as discussed in Methods. In the configuration of the baseline model, total number of trees was set to 100 while the maximum depth of each tree was 10. A 10:1 weight ratio was assigned to true positive and false positive labels, respectively, and information gain was used as the criterion to split the nodes in each tree. The baseline model achieved 100% accuracy on the training data, and 0.98 F-score in a ten-fold cross-validation.

### Finding Optimal Three-class Model

Fine-tuning parameters of the classifier was performed using the validation set, which consisted of 2,329 SNVs from 66 somatic tumor samples. Example parameters evaluated in this step included weight ratio between the true positives and artifacts, maximum depth of a tree and total number of trees in the forest. Due to the characteristics of data available and the intrinsic behaviors of the computational models, a classifier was often more confident about some predictions than others presented with new data. The prediction intervals in the range of [0, 1] were used to measure the level of confidence *(37)*. Specifically, a value equal or close to one indicated the classifier was confident the variant was real while a value equal or close to zero indicated the classifier was confident the variant was an artifact. Therefore, the third class of “uncertain” variants was defined as less confident classifications inferred by the prediction intervals. The ideal model should yield minimized number of misclassifications while the prediction intervals of the misclassifications should be far away from the two ends of the range.

Guided by these heuristics, the candidate classifier instances were sorted by the number of misclassifications and then the difference between the highest and lowest prediction intervals among the misclassifications. The optimal classifier was identified, along with the boundaries of the third class defined by the prediction interval: [0.05 - 0.9]. The prediction intervals of all of the misclassifications were within the range, with sufficient margins to the boundaries. The resulting RF consisted of 51 trees whose maximum depth was 10 and the weight ratio between the two classes was 101:1. Using this classifier, SNVs with prediction intervals in the range of [0 - 0.05) were labeled as non-reportable, those with prediction intervals in the range of (0.9 - 1] were labeled as true while the rest were labeled as “uncertain” and hence require manual inspections (Figure 4). In the validation set, 496 out of 526 (94.3%) TP SNVs were predicted real, 1779 out of 1803 artifacts (98.7%) were labeled false while the remaining 54 variants (2.3%) were labeled “uncertain”.

**Figure 4.**
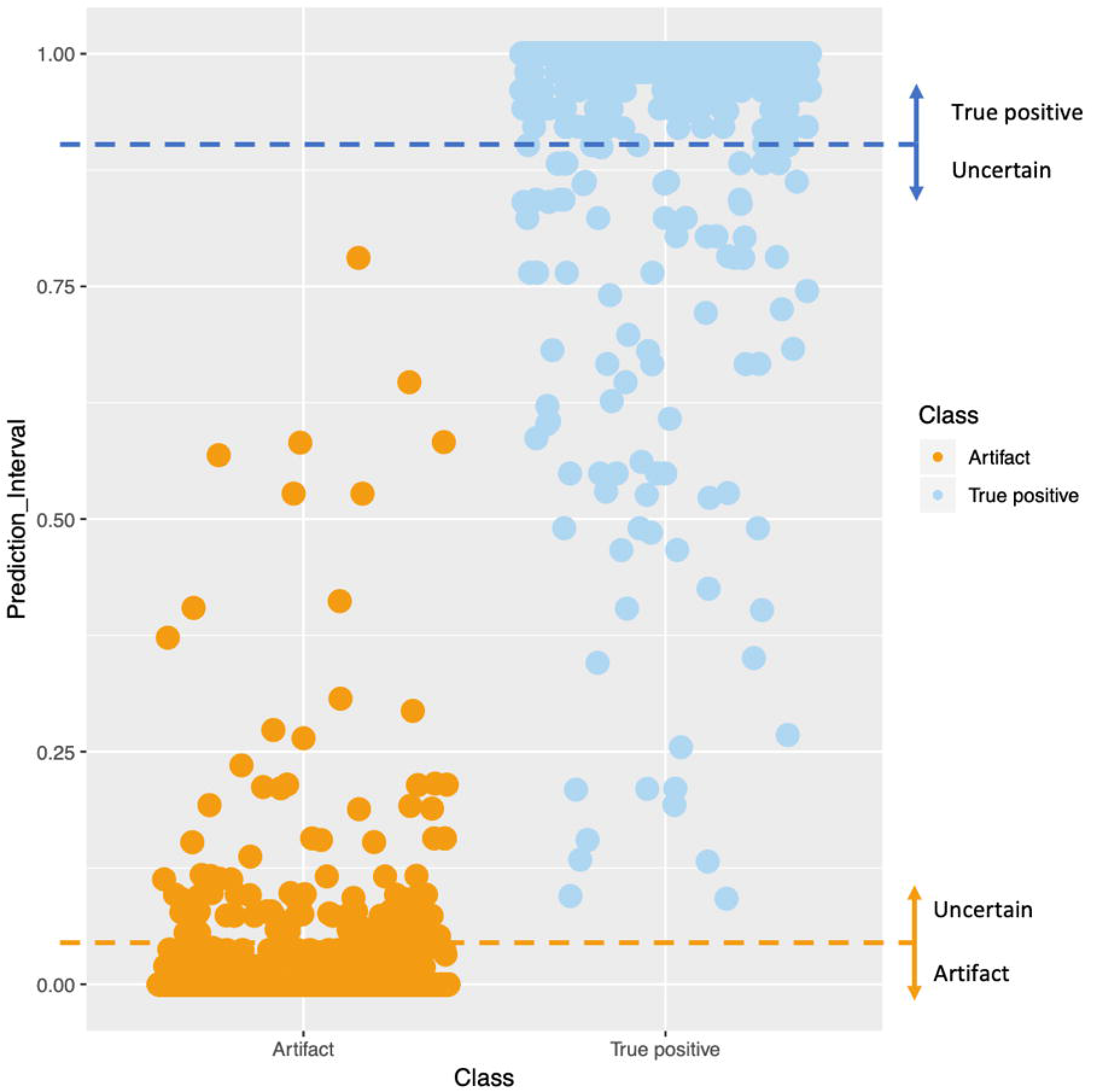
Plot of prediction intervals of variants in the test set including the thresholds defining the class of “uncertain”. 192 variants of prediction intervals between 0.05 and 0.9 were assigned the label of “uncertain”.

### Benchmarking Using Test Set

We then further evaluated the optimal classifier on the independent test set, which consisted of 5,587 SNVs from 141 somatic tumor samples. Applying the classifier, 1252 out of 1341 (93.3%) TP SNVs were predicted real, 4143 out of 4246 (97.6%) artifacts were labeled artifacts while the remaining 192 (3.4%) SNVs were labeled as “uncertain”. More importantly, none of the TP SNVs were misclassified as sequencing artifacts or *vice versa*, while only 3.4% of the SNVs did not receive a definitive label and required further manual investigation.

### Feature Importance and Uncertain SNVs

To determine the relative contribution of each feature in the computational model, feature importance analysis was performed on the optimal classifier. The importance of a feature was measured by the decrease in accuracy of the classifier when the values of the feature were randomly permutated. Strand bias was recognized as the most important feature, followed by alternate allele frequency and batch effect. The complete list of feature importance is summarized in Table 1. Additionally, further investigation did not suggest strong pairwise correlations among features. More details are presented in the Supplemental Methods.

**Table 1.** Feature importance

| Feature | Importance |
| --- | --- |
| Strand Bias | 0.38 |
| VAF | 0.21 |
| Batch effect | 0.17 |
| Alternate Coverage | 0.11 |
| Dissimilarity to normal sample 2 | 0.09 |
| Dissimilarity to normal sample 1 | 0.03 |

The optimal classifier agreed with the manual inspection in 97.7% (2275/2329) and 96.6% (5395/5587) of calls in the validation and test sets, respectively. However, 54 (2.3%) and 192 (3.4%) variants from respective sets were labeled “uncertain” and hence discordant from their original labels. Among these 246 “uncertain” variants, 119 were real mutation events while the rest 127 were manually rejected by the genome scientists. These discrepancies would not impact clinical outcome as these variants would be manually inspected.

### Impact on Clinical Workflow

In order to assess impact of using of implementing the optimal model in clinical variant review process in the lab, we measured combined hands-on time of first and second variant review steps for 203 cases before and for 211 cases after implementation. Average hands-on time before implementation was 240 minutes compared to 89 minutes after the model was implemented which is an improvement of 63%. This has enabled the lab to reduce the turnaround time and increase variant review capacity by 42% with the same number of genome scientists.

## Discussion

Manual inspections on the variants detected from tumors are commonly implemented in clinical laboratories for quality control. This will delay the release of results on which oncologists rely to deliver timely treatments for their patients. In order to automate the process, we developed a computational classifier to distinguish sequencing artifacts from the true positive events. We limited our study to SNVs of non-FFPE samples because they constituted the majority of the variants detected and there was an insufficient number of indels to train computational models. As a result, the optimal RF-based three-class model demonstrated high accuracy and utility on the validation set and the independent test set. Overall, 96.6% of the SNVs received a definitive label and hence be exempt from the manual screening process.

To better understand features influencing the 246 variants labeled “uncertain” by the optimal model in the validation and test sets, they were further examined and plausible explanations were as follows. For 115 variants, the coverage might be too low for the computational model to confidently determine their validity (average coverage <50x). Another 65 “uncertain” variants might suffer from low VAF (VAF <10%), despite the overall coverage at the locus. The rest were labeled “uncertain” likely because of their complex feature manifestations. For example, a TP variant labeled “uncertain” exhibited high strand bias for it resided in a GC rich region. Another “uncertain” variant of similar characteristics was, however, an artifact, where its lowered similarity to normal controls was primarily due to low read depths on the control samples. In these cases, variants were warranted to be manually inspected and subject to confirmatory methods to determine its validity.

Compared to the recent study which generated over 70 features for each variant, our dataset did not include many of those features such as tumor type or average number of mismatches in the aligned reads *(20)*. While those characteristics could be helpful in refining the sequencing data in the abovementioned study, we decided they were less applicable to the clinical lab practice. Some of those characteristics were correlated with our selected features and hence redundant to the machine learning models. Meanwhile, the minimal set of highly pertinent features we selected was a close reflection of the manual screening procedure. Our study suggests it is possible to achieve similar or better performance in pediatric tumor sequencing using limited yet carefully designed features compared to the abovementioned study using adult sequencing data.

There are several limitations to this study. All of the data used in the study was generated from the cancer genomic diagnostic laboratory at CHOP, hence, it is possible that the model could be partial to the latent characteristics that are laboratory-specific. However, the robustness demonstrated in the results indicated the methodology is applicable to overall variant screening carried out in other laboratories.

Additionally, the machine learning models were trained and optimized using data from manual review. Ideally, all variants used for training the models need to be confirmed by orthogonal methods but this would not be feasible for the reasons explained in Introduction. Therefore, it was possible that noise might have been introduced during the manual label process. For instance, the majority of the polymorphic variants were labeled as “real” while some others were labeled “non-reportable”. Such variants were difficult to be consistently classified as polymorphism or sequencing artifact without further confirmation. Although they did not present any clinical significance, these variants might have contributed imperfections to the models. Consequently, the classifier was less confident on these variants and labeled most of which “uncertain”.

Our classifier did not aim to distinguish somatic and germline variants in cancers. The majority of the clinical laboratories in the US offer tumor-only assays, due to the clinical challenges of obtaining normal specimen such as specimen adequacy concerns in pediatric patients, the logistics of acquiring normal samples, and the requirements for complex consent forms *(38)*. Specimen used in this study were submitted for tumor-only tests, but might contain a certain percentage of germline tissues. Thus, inherent challenges remain in distinguishing germline variants based on tumor-only tests, which make it difficult to generate confident training data to build the model to classify the germline variants*(38)*. In future studies similar approaches could be applied to identify germline variants with minor adjustment provided well curated data is available.

In summary, sequencing artifacts caused by a number of fundamentally persistent issues could overwhelm *bona fide* variants in somatic tumor sequencing. Manual screening of the variants is subjective and labor intensive. Here, we have presented an approach to apply machine learning methods to systemically identify TP SNVs from artifacts in pediatric non-FFPE tumors. We have shown the accuracy and robustness of our approach, as well as the reduced bias and gained efficiency implementing the model in a clinical setting.

## Acknowledgements

We would like to thank Sarah Lipson from Wake Forest University for her contributions in editing and improving the quality of the manuscript.

