## Supplementary Information for "Using Machine Learning to Identify True Somatic Variants from Next-Generation Sequencing"

**Supplemental Methods**

**Feature correlations**

potential correlation between features could make it challenging to interpret relative feature importance in tree-based classifiers. Therefore, we also investigated the correlations between the features. Values of the features for each variant were normalized to the range of 0 to 1 before Spearman test was carried out to evaluate the pair-wise correlations. The results did not reveal any strong correlations between the features which demonstrated robustness of our features (Figure S2).
