## Supplementary figures and images for "Using Machine Learning to Identify True Somatic Variants from Next-Generation Sequencing"

### Fig_S1_breakdown.tiff

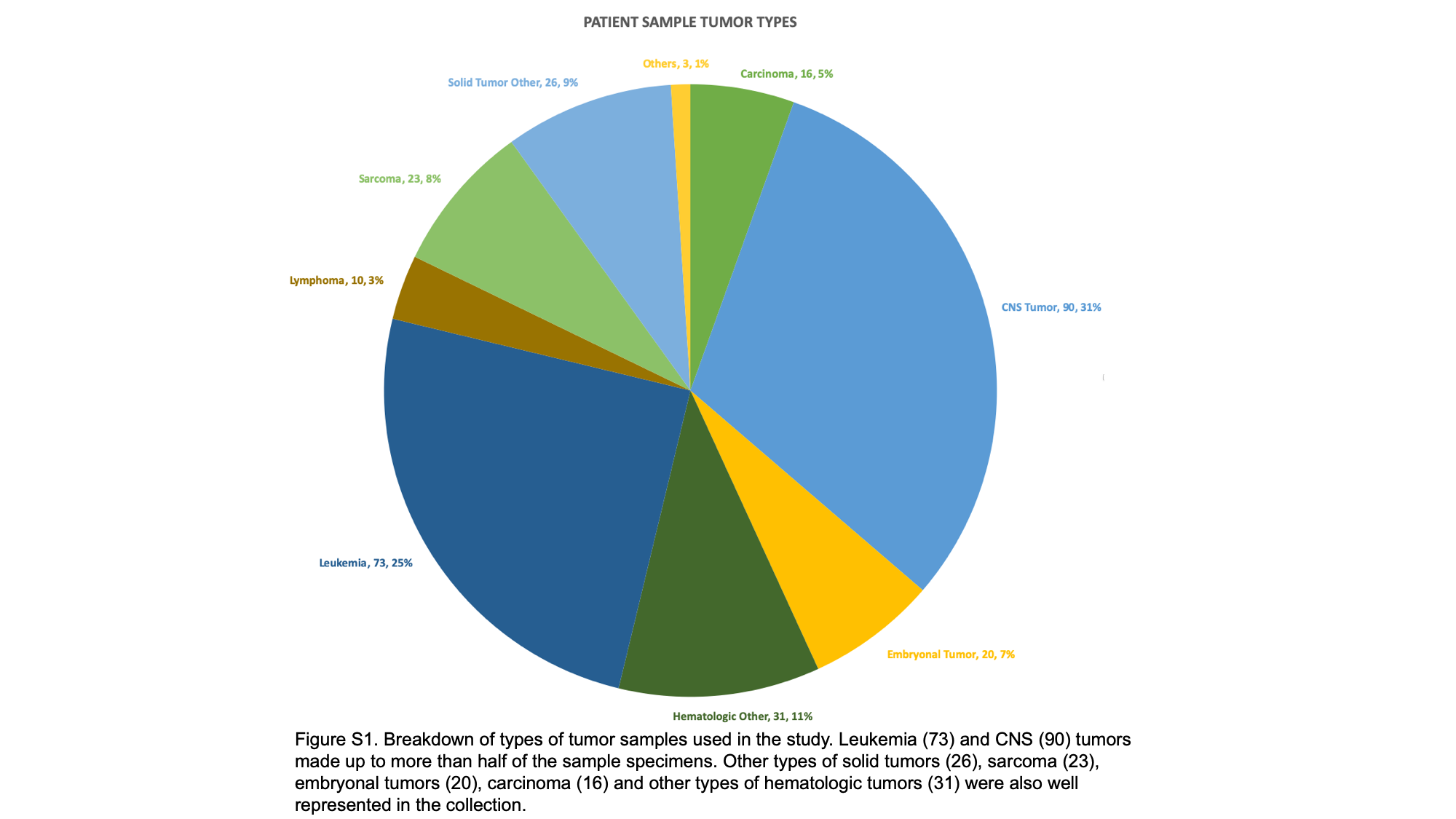

### Fig_S2_correlations.tiff

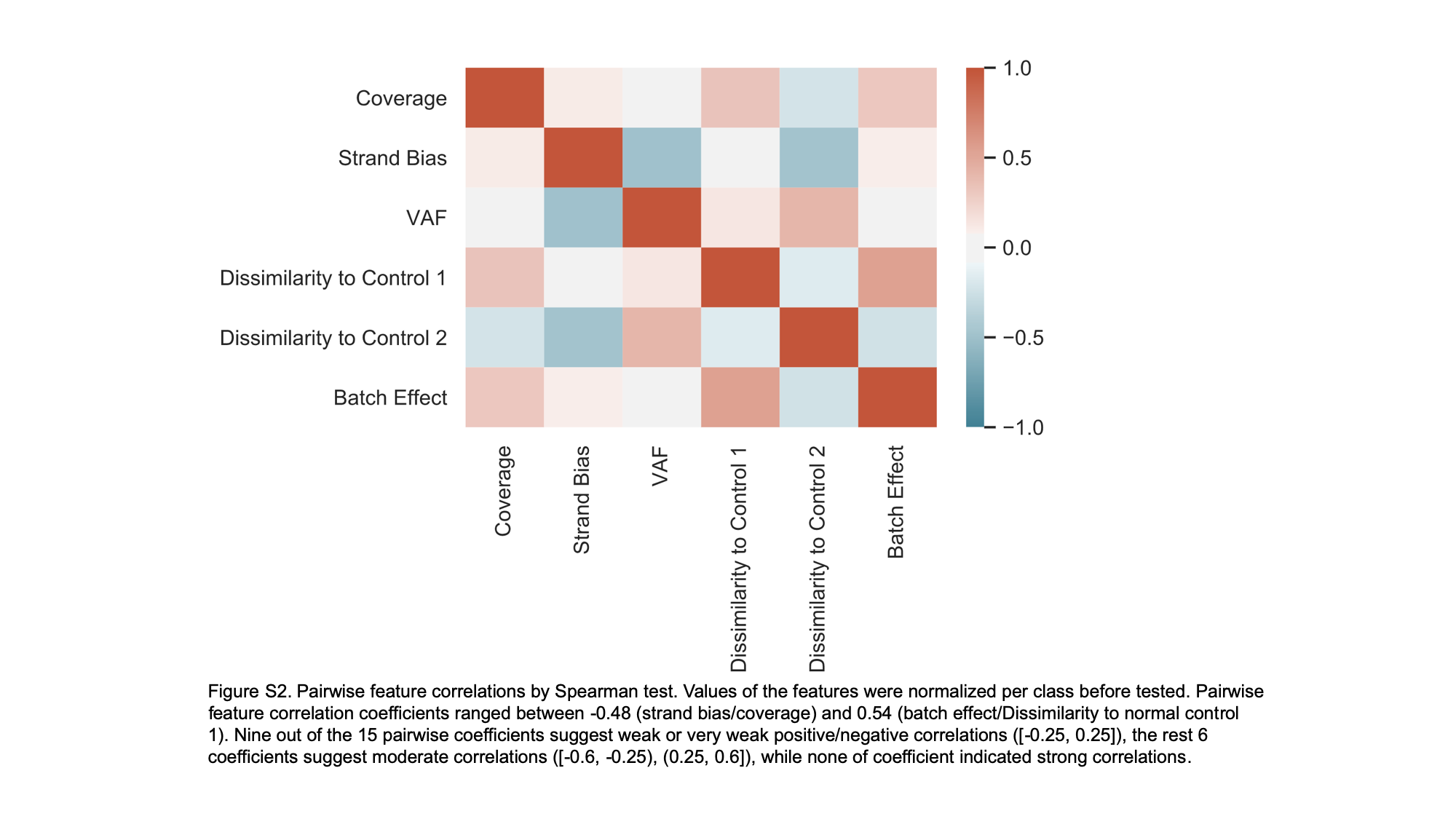
